# 3-D deconvolution of human skin immune architecture with Multiplex Annotated Tissue Imaging System (MANTIS)

**DOI:** 10.1101/2023.01.13.523748

**Authors:** Manon Scholaert, Raissa Houmadi, Jeremy Martin, Nadine Serhan, Marie Tauber, Emilie Braun, Lilian Basso, Eric Merle, Pascal Descargues, Manuelle Viguier, Cécile Lesort, Benoît Chaput, Jean Kanitakis, Denis Jullien, Cristina Bulai Livideanu, Laurence Lamant, Emeline Pagès, Nicolas Gaudenzio

## Abstract

Routine clinical assays, such as conventional immunohistochemistry, often fail to resolve the regional heterogeneity of complex inflammatory skin conditions. Here we introduce MANTIS (Multiplexed Annotated Tissue Imaging System), a flexible analytic pipeline compatible with routine practice, specifically-designed for spatially-resolved immune phenotyping of the skin in experimental or clinical samples. Based on phenotype attribution matrices coupled to α-shape algorithms, MANTIS projects a representative digital immune landscape, while enabling automated detection of major inflammatory clusters and concomitant single-cell data quantification of biomarkers. We observed that severe pathological lesions from systemic lupus erythematosus, Kawasaki syndrome or COVID-19-associated skin manifestations share common quantitative immune features, while displaying a non-random distribution of cells with the formation of disease-specific dermal immune structures. Given its accuracy and flexibility, MANTIS is designed to solve the spatial organization of complex immune environments to better apprehend the pathophysiology of skin manifestations.

## Introduction

The skin acts as a barrier organ that separates the body from the external environment. Upon inflammation, blood-circulating immune cells are recruited to help orchestrate the cutaneous immunity and are often nested nearby key structural elements (e.g., postcapillary venules, hair follicles, dermal-epidermal junction, etc.)^1,2^. In pathological settings, the nature and activation status of the skin immune landscape often represent precious biological information that can help establish an accurate diagnosis, apprehend interpatient heterogeneity and select the most appropriate treatment. The use of imaging-based approaches to identify cutaneous immune cells is still challenging because of the high level of autofluorescence arising from the tissue itself, the potential spectral spillover when more than 4 fluorochromes are used simultaneously and the entanglement of all cells within thick and polarized structural appendages.

The vast majority of microscopic diagnoses of inflammatory skin conditions relies on repeated immunohistochemistry (IHC) analysis of one or two proteins and/or hematoxylin eosin (H&E) staining in thin (2 to 5 μm) formalin-fixed, paraffin-embedded (FFPE) specimens^3,4^. While such two-dimensional approaches are reproducible and suitable for routine practice, they do not permit to apprehend the complex topology and heterogeneity of immune cells^5^, in particular those nested in-between epidermal appendices. The development of image-based histo-cytometry, which consists in analyzing segmented multicolor images with classical flow cytometry gating strategies, has paved the way toward the development of sophisticated image-generation systems coupled to computational imaging^6^. Recently, highly-multiplexed imaging systems have significantly advanced our understanding of tissue-resident immune subsets and of their spatial distribution with regards to tissue structures, with a strong focus on cancer samples and tumor heterogeneity, such as CODEX^7,8^, MIBI-TOF^9^, IMC^10^, MuSIC^11^, CyCIF^12^, Cell Dive^13^ and others^14^. While multiplexed imaging has an immense potential, there is a strong need to democratize these methods with the use of inexpensive instrumentation compatible with standard tissue processing and coupled to an analysis interface that is user-friendly enough to be used in routine practice.

Here we present an integrated framework primarily designed for spatially-resolved immune cells phenotyping in FFPE human skin biopsies. We first set up a simple and inexpensive method to acquire 10 fluorescent signals simultaneously and in 3-D using a classical confocal microscope. We next designed MANTIS (Multiplexed Annotated Tissue Imaging System), an adaptable and interactive analytical system which automatically generates a digitalized version of the skin immune landscape and enables single-cell quantitative data visualization. Based on these settings, MANTIS could be implemented in most laboratories coupled to existing confocal equipment to bridge the gap between sophisticated research tools and standard-of-care diagnostic procedures with minimal human intervention.

## Results

### Extraction of single-cell statistics from skin sections by combining conventional confocal laser-scanning microscopy and computational imaging

We developed a simple method to generate 3-D multiplexed fluorescent images from FFPE (20 μm thick) skin biopsies which could be implemented in most research or clinical laboratories on existing equipment. Skin sections were first stained with different panels of commercially-available fluorochrome-coupled antibodies added simultaneously, then quenched to avoid excessive natural autofluorescence of skin structural elements (**Fig. 1A**). We acquired 3-D fluorescent multiplexed images with a conventional inverted confocal laser-scanning microscope equipped with 5 laser lines, 5 detectors and a 40X oil immersion objective, using a strategy of sequential acquisition composed of fast consecutive steps (**Fig. 1B**; the detailed description of optical paths and lasers of our 8-year-old Leica SP8 system is provided in the Materials and Methods section). This setting enabled the acquisition of 8 to 10 fluorescent channels, on a system primarily designed for 4 colors, over a skin section of the following 3 dimensions, 0.6 mm (x) x 0.4 mm (y) x 20 μm (z) within 25 minutes. The obtained 3-D images were then deconvoluted and compensated to correct 3-D fluorescent spectral spillovers (**Fig. 1C, D**) using the Huygens software (Scientific Volume Imaging), a strategy routinely applied in flow cytometry to combine multiple fluorochromes simultaneously^6,15^. Compared to classical segmentation strategies based on nucleus expansion^16^, which often lead to under or over-estimation of cellular cluster composition, we used the general immune biomarker CD45 as a robust immune staining visualized in most skin-resident immune cells to constitute the core of our cell segmentation strategy for future single immune cell statistics extraction (**Fig. 1E, F**). Using the Isosurface algorithm of the Imaris software (Bitplane) we next modeled the 3D fluorescence signal of CD45 for individual immune cells and exported a corresponding single-cell database composed of the mean fluorescence intensity (MFI) of all individual biomarkers and precise x, y, z tissue coordinates obtained with a resolution of 299 × 299 × 999 nm/voxel (**Fig. 1F**). We found that CD45-based segmentation enabled an efficient isolation of single immune cell characteristics, even when those were found aggregated around dermal structural elements. Overall, we demonstrate that it is possible to extract a 10 parameter-single-cell database using regular confocal equipment coupled to basic computational imaging steps.

**Figure 1.**
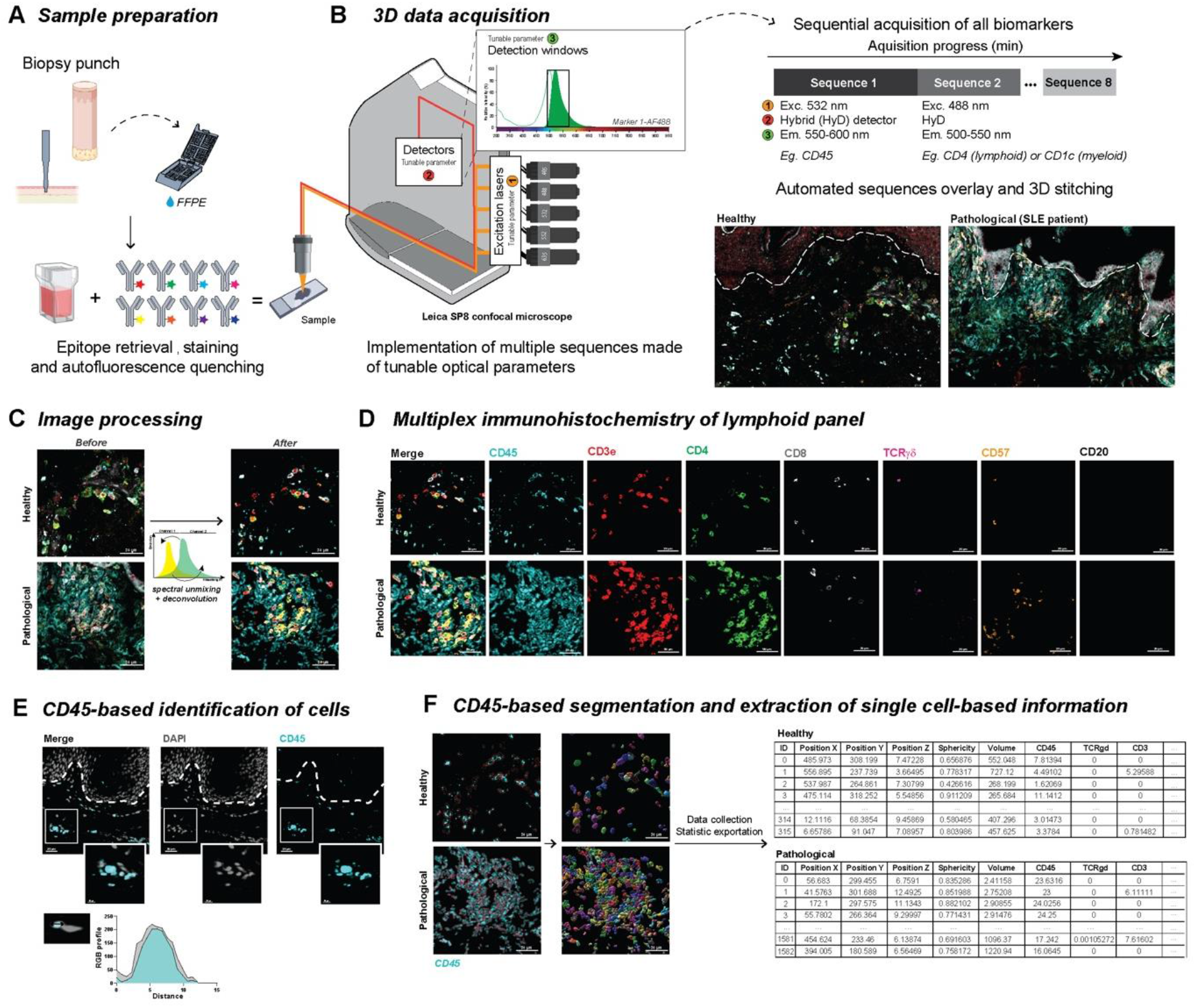
Between-stack microscope configuration allows sequential acquisition of 7+ channels with classical image processing. **A**, Sample preparation. FFPE-skin sections were cut and stained for myeloid and lymphoid panels after appropriate epitope retrieval and autofluorescence quenching. Sample images were then acquired using a SP8 confocal microscope from Leica Microsystems as described in B. **B,** Microscope configuration and acquisition settings. Mosaic sequential images were acquired using the between-stack configuration with tunable detection windows. Sequences were overlaid and 3-D-stitched. An example of data acquisition is given for healthy (left panel) and pathological (lupus erythematosus [SLE], right panel) skin. **C**, Deconvolution of regions of interest and spectral unmixing. Acquired 3-D images were deconvoluted and compensated to correct optical aberrations and 3-D fluorescent spectral spillovers. **D,** Representative 3D multiplex image of healthy (upper panel) and pathological SLE (lower panel) skin sample for lymphoid panel, staining CD45, CD3, CD4, CD8, TCRγδ, CD57 and CD20. **E,** Co-localization of DAPI and CD45 staining and respective RGB profiles. **F**, Segmentation and single-cell database creation. Cell segmentation using the CD45 fluorescence channel allowed efficient isolation of individual objects, i.e., immune cells. Individual object statistics (*xyz* coordinates, sphericity, volume and Mean Fluorescence Intensity) were extracted for each sample.

### Analysis of the skin immune landscape using MANTIS phenotypes attribution matrices

Based on the literature, we designed two panels composed of antibodies directed against immune biomarkers suitable to generate a non-exhaustive overview of lymphoid and myeloid cell landscape of the skin, with an average cost of approximately $65 per sample. The combination of CD45, CD3e, CD4, CD8, TCRγδ, CD20 and CD57 (a terminally sulfated glycan carbohydrate epitope shared by NK and T cells with high cytotoxic potential^17,18^) allows to identify the following lymphoid cells: conventional CD4 and CD8 T cells (being CD57^low^ or CD57^high^), CD4^+^ CD8^+^ double positive (dp) T cells^19^, CD4^-^ CD8^-^ double negative (dn) T cells^20^, γδ T cells, B cells and NK cells (**Table 1**). The combination of CD45, CD207, CD1c, HLA-DR, CD123, Siglec-8, myeloperoxidase (MPO) and tryptase allows to identify the following myeloid cells: Langerhans cells, Langerin^+^ (CD207^+^) dermal dendritic cells (dDCs) and Langerin^-^ dDCs, eosinophils, basophils, neutrophils and mast cells (**Table 1**). The activation status of DC, Langerin^+^ DC and LC was investigated using levels of HLA-DR expression. A detailed list of excitation/emission/detection strategies is provided in **Table 2**.

We next aimed to develop an adaptable analytical system that could integrate and batch-process extracted single-cell databases and enable an unsupervised phenotyping of immune subsets. To address this latter challenge, we developed MANTIS, an interactive digital tool based on phenotype attribution matrices inspired by the analytical logic of single-cell RNA sequencing that identifies correlations between the single-cell database and the expression profiles of different cell types (**Fig. 2A**). Such an analysis is possible by computing Spearman’s Rho correlation, which accommodates non-linear relationships in the expression values (i.e., in our case the collected MFI of each biomarker). In practice, MANTIS runs instantaneously a pairwise Spearman’s correlation analysis, for each detected single-cell, against selected combinations of biomarkers to identify the immune subsets annotated in the phenotype attribution matrices. The output information is the attribution of specific Rho values per single cell which then automatically finds the best match of cellular identity and generates associated quantitative statistics (**Fig. 2A-C, Supp Fig. 1A, B**). As a proof of concept, we generated data from two serial sections of an acral lesion from a patient with systemic lupus erythematosus (SLE, i.e., lupus chilblains) stained with a lymphoid and a myeloid panel. The fast 3-D acquisition of one region of interest (ROI) composed of 6 fields of view (i.e., 0.6 mm (x) x 0.4 mm (y) x 20 μm (z)) enabled the annotation of 519 myeloid cells and 708 lymphoid cells for a total of 19 different immune subsets identified (**Supp Fig. 1C**, **Fig. 2D**). One can then decide to visualize annotated immune populations using either a heatmap, in which the MFI of individual biomarkers is displayed per single cell (**Supp Fig. 1D**), or a graph-based dimensionality reduction, i.e., t-distributed stochastic neighbor embedding (t-SNE), specifically designed for visualizing clusters of populations and corresponding expression of biomarkers per cluster (**Fig. 2E**).

**Figure 2.**
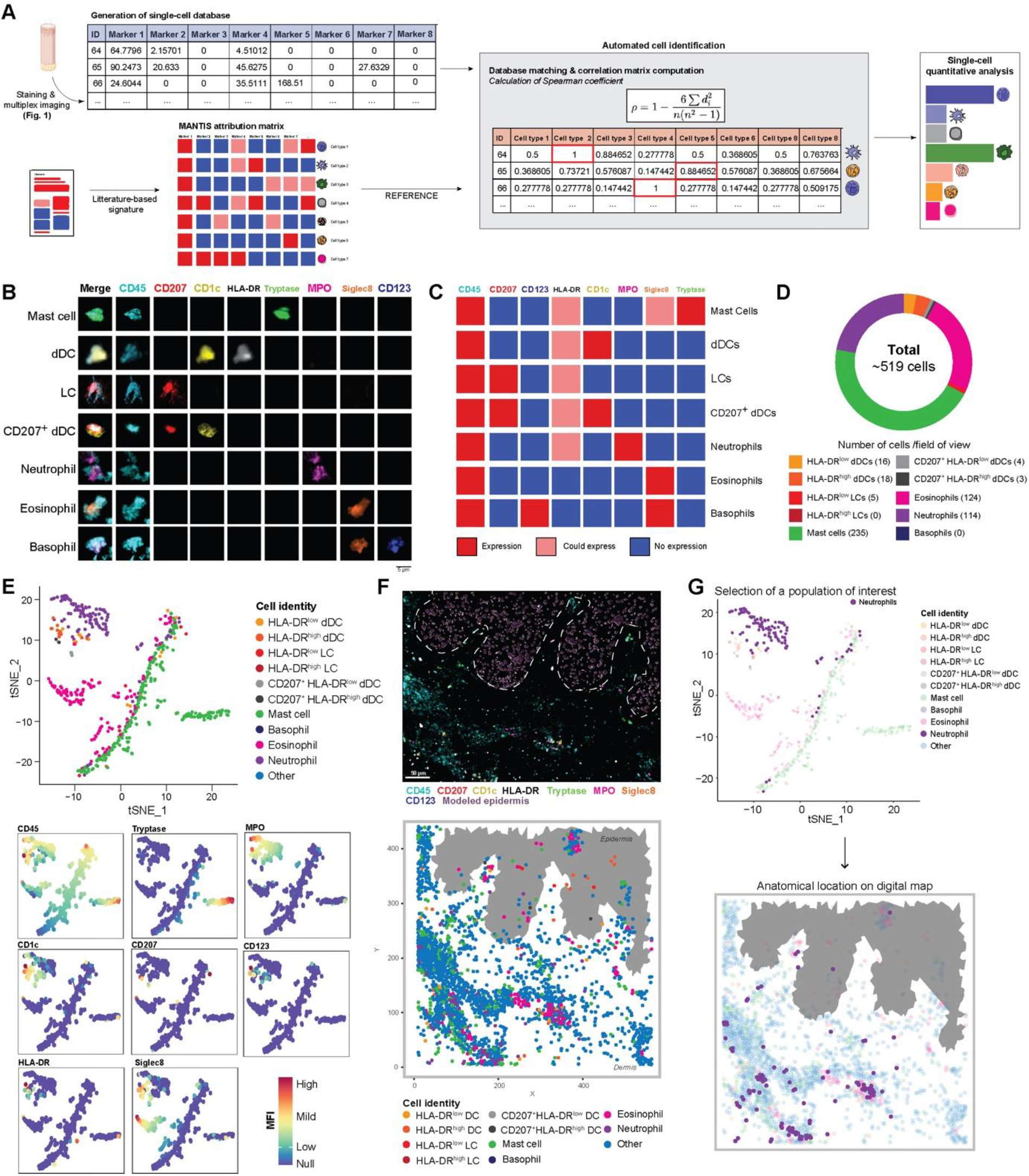
MANTIS algorithm allows automated cell type attribution and interactive exploration of skin myeloid immune topology. **A,** Automated tissue annotation. A reference attribution matrix defining the literature-based theoretical signature of a particular cell type was constructed and designated as MANTIS attribution matrix. A correlation matrix calculating Spearman coefficient between the single-cell database and MANTIS attribution matrix was computed. Each segmented cell was annotated to the cell type having the highest correlation coefficient, and cell type proportions were extracted. **B**, Single-cell staining of all used biomarkers in identified myeloid cells. **C**, MANTIS simplified attribution matrix for myeloid panel. **D**, Tissue annotation and cell proportion of pathological (SLE) skin. **E**, Representative t-SNE plot of myeloid cell populations (upper panel) and MFI levels of used markers (colored intensity scale, lower panel). **F,** Representative 3D confocal multiplex image (upper panel) and associated digital map (lower panel) of pre-designed MANTIS myeloid panel of pathological (SLE) skin. **G**, Interactive reverse-gating. A population of interest (neutrophils) was selected on the tSNE plot. Recomputation of the corresponding digital map enabled the visualization of the anatomical distribution of this particular population in the skin biopsy.

A particularly challenging aspect of multiplexed imaging technologies is to circumvent the spatial distribution of immune cells with respect to longitudinal and polarized structural elements (e.g., epidermal appendages of the skin) in thick tissue sections. We developed an interactive software interface that contextualizes the immune topology of the skin by replacing all annotated single immune cells within their 3-D spatial context and leverages the natural autofluorescence of keratinocytes to model the epidermal layer to facilitate biopsy orientation (**Fig. 2F, Supp Fig. 1E, F**). The algorithm allows to use two complementary analytical approaches and to switch from one to the other with a simple drawing tool (**Supp Video 1**). The analysis can start from the visualization of the skin digital immune landscape, then be pursued with the investigation of the immune composition of defined microregions via instantaneous re-computation of drawn ROI. Conversely, it is also possible to start from all annotated immune cells on a t-SNE graph, draw around subsets of interest, and immediately visualize their anatomical distribution in the skin digital immune landscape (**Supp Video 1, Fig. 2G**). Taken together, these data suggest that MANTIS interactive analytical system can be used to compute the 3-D spatial organization of immune and structural elements of inflammatory skin samples from patients.

### Quantitative validation of MANTIS annotation system using healthy-looking skin and inflammatory pathological lesions

With the constant increase in the number of cases, a large panel of putative skin manifestations of COVID-19 have been observed worldwide^21,22^, including an unprecedented high rate of acral lesions which represent ~ 75% of all cases and commonly named “COVID-Toes”^23–26^. Such manifestations (**Supp Fig. 2A**), compared to non-inflamed healthy-looking skin (**Supp Fig. 2B**), are associated with an important immune cell infiltration (**Supp Fig. 2C**) and tend to develop in young patients with no or very mild respiratory symptoms^26,27^. While some pathological features of those lesions have been described ^28,3,29^, a precise analysis of their spatial immune profile is currently missing, which impairs the development of a clear readout to better diagnose and treat these rare cutaneous lesions. A possible explanation could be a collateral clinical manifestation of an efficient anti-viral type 1 interferon response, since acral lesions are also commonly observed in patients with interferonopathies, such as the Aicardi-Goutières syndrome ^30^ ref and SLE^31^. With this in mind, we decided to benchmark the effective performance of MANTIS to resolve the immune topology of skin lesions of similar clinical severity from 5 patients with COVID-toes, 2 patients with the multi-system inflammatory syndrome (MIS), which is clinically similar to Kawasaki syndrome (i.e., a rare severe systemic inflammatory condition triggered by SARS-CoV-2 infection, named there after “Kawasaki syndrome”) and 3 patients with SLE chilblains. Abdominal skin biopsies from 5 healthy-looking controls were used to set the baseline of a natural steady-state immune environment, albeit from a distant anatomical region.

We validated the quantitative performance of MANTIS to annotate immune cells by calculations of statistical correlations with a supervised approach of histo-cytometry^15,32^ applied on the same datasets for each antibody panel in all skin samples. This last method consists in a manual gating of immune subsets on the same principle used in traditional flow cytometry. A total of 20,464 single CD45^+^ immune cells were identified with the following distribution per condition: 1,670 immune cells in 5 healthy-looking skin samples (i.e., with 895 myeloid and 775 lymphoid cells), 1,703 in 2 Kawasaki syndrome patients (i.e., with 932 myeloid and 775 lymphoid cells), 5,076 in 3 SLE chilblain patients (i.e., with 1,560 myeloid and 3,516 lymphoid cells) and 12,015 in 5 COVID-toes patients (i.e., with 2,838 myeloid and 9,177 lymphoid cells). A classical gating strategy based on mutually exclusive biomarkers was used to assess the presence of myeloid (**Supp Fig. 3A**) and lymphoid (**Supp Fig. 4A**) cell subsets by histo-cytometry. We identified a total of 19 different immune subsets and found very similar distributions of cell counts by either supervised histo-cytometry or unsupervised MANTIS algorithm (**Supp Fig. 3B-E, 4B-E**). The calculated R coefficients were between 0.75 and 1, regardless of the antibody panel, the patients analyzed or the disease (**Supp Fig. 3F** and **4F**). Importantly, we observed that all healthy-looking skin samples exhibited a proportion of immune cells aligned with previously described skin-resident immune populations at steady state in human^1,33^. However, we noted a slightly higher tendency to detect rare populations of blood-circulating CD45^+^CD3^-^CD20^+^ B cells or CD45^+^HLA-DR^-^CD123^+^Siglec8^+^ basophils, only when 3-D images were computationally analyzed with MANTIS (**Supp Fig. 3B** and **4B**). This is consistent with the fact that the skin is a highly-vascularized tissue and that recent studies identified rare B cells in healthy skin^34^.

Having validated the quantitative and qualitative performance of MANTIS-based annotation, we next defined a high-level view of the complex immune environment of pathological lesions from all 10 patients. Compared to healthy-looking samples, pathological lesions contained large immune infiltrates confirming their inflammatory status (**Fig. 3A, B**). All three conditions were associated with an infiltration of myeloid cells composed of a large number of neutrophils, eosinophils, mast cells and conventional CD45^+^CD1c^+^CD207^-^HLA-DR^+^ dDCs (**Fig. 3A, C**). While detected in relatively low numbers in all analyzed skin samples, no difference was observed between healthy-looking and pathological samples for CD45^+^CD1c^-^CD207^+^HLA-DR^+^ LCs or CD45^+^CD1c^+^CD207^+^HLA-DR^+^ dDCs populations.

**Figure 3.**
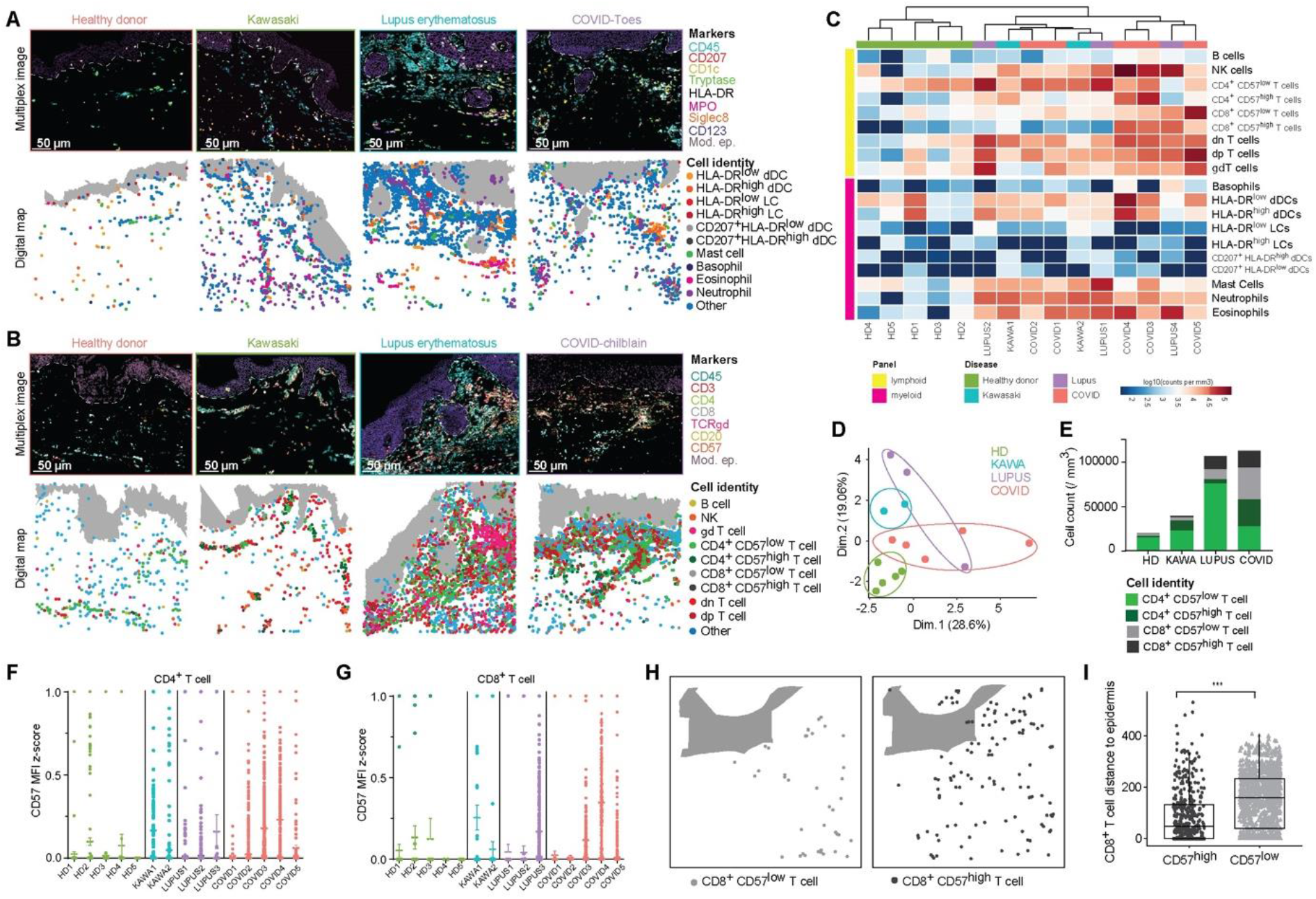
3-D quantitative and spatial analysis of skin immune cells at the cellular level provide insight into disease signatures. **A, B,** Representative 3-D confocal multiplex images (upper panel) and associated digital maps (lower panel) of pre-designed MANTIS myeloid (A) and lymphoid (B) panels of healthy and pathological skin. **C**, Representative heatmap of lymphoid and myeloid cell densities in logarithmic scale with hierarchical clustering. **D**, Principal Component Analysis (PCA) of immune signatures of healthy and diseased skin. **E**, Cell count per mm^3^ of CD57^low^ and CD57^high^ T cells. **F, G**, Dotplot of CD57 Mean Fluorescence Intensity (MFI) z-score in CD4^+^ (F) and CD8^+^ (G) T cells in healthy and diseased skin. **H, I**, Representative digital map (H) and mean distance to epidermis (in μm [I]) of CD8^+^ CD57^low^ (left panel) and CD57^high^ (right panel) T cells in COVID skin lesions.

Compared to Kawasaki syndrome patients, SLE and COVID-19 patients tended to have an increased proportion of lymphoid cells (**Fig. 3B, C**), with an enrichment in conventional CD4^+^ or CD8^+^ T cells and NK cells, and to a lesser extent in γδ T cells. Interestingly, we also observed double-positive (dp) CD45^+^CD4^+^CD8^+^CD3^+^TCRγδ^-^ and double-negative (dn) CD45^+^CD4^-^ CD8^-^CD3^+^TCRγδ^-^ T cells in all inflamed and some healthy-looking samples, albeit in smaller numbers (**Fig. 3B, C** and **Supp Fig. 3**). Such populations of T cells were often understudied, as CD4 and CD8 biomarkers are thought to be mutually exclusive, however they have been often reported in autoimmune and chronic inflammatory disorders^19,35^, including SLE^36,37^.

We next performed an unsupervised clustering of all patients and healthy-looking controls based on the quantitative analysis of their immune signature using both a detailed heatmap based on immune profiles (**Fig. 3C**) and a principal component analysis (PCA) per patient (**Fig. 3D**). Healthy-looking skin samples clustered together, with no apparent relationship with the pathological samples (**Fig. 3C, D**). Kawasaki syndrome and COVID-toes patients had a tendency to form disease-specific clusters, while SLE patients were distributed between both conditions (**Fig. 3C, D**). Even though these data were obtained on a restricted number of patients, they suggest that all analyzed pathological lesions displayed common quantitative immune features (**Fig. 3C**), with nevertheless potential disease-intrinsic characteristics suggested upon analysis with a dimensional reduction PCA (**Fig. 3D**). To explore further this hypothesis, we refined our analysis by investigating the activation status of conventional CD4^+^ and CD8^+^ T cells based on their expression level of CD57, a biomarker classically associated with a high cytotoxic potential (i.e., pro-tissue damage) during viral infections and autoimmune disorders, including COVID-19^38^. We found that, compared to other pathological conditions, 3 COVID-toes cases were particularly enriched in CD4^+^ and CD8^+^ T cells exhibiting high levels of CD57 (i.e., CD57^high^; calculated as CD57 MFI z-score, **Fig. 3E-G**).

During inflammatory skin conditions, cytotoxic immune cells can relocate nearby to/in contact with keratinocytes and contribute to severe epidermal damage^39,40^. In order to calculate the anatomical location of all immune cells with respect to the epidermal layers, we acquired the spatial coordinates of the modeled epidermis. We next incorporated into MANTIS a k-dimensional tree algorithm^41,42^, which automatically decomposes the structural element coordinates (i.e., as exemplified here with the epidermis) into virtual subspaces and enables to calculate the nearest neighbor to each immune cell (**Supp Fig. 5A**). A batch calculation of the distance of each individual cell can then be visualized under the format of a heatmap, providing a quick overview of the dataset (**Supp Fig. 5B**). We found that HLA^high^ dDCs (**Supp Fig. 5C**), NK cells (**Supp Fig. 5D**) and CD8^+^ CD57^high^ T cells (**Supp Fig. 5E**) were all significantly enriched near the epidermal layer in cases of COVID-toes. Importantly, CD8^+^ CD57^low^ T cells were not found enriched in the epidermis (**Fig. 3 H-K**), suggesting a biological link between the expression levels of CD57 and epidermal migration in CD8 T cells. Even though the number of patients studied is limited, these findings strongly suggest the potential formation of tissue damaging subepidermal inflammatory clusters composed of cytotoxic T cells and NK cells in COVID-toes.

### MANTIS enables topographic exploration of skin lesions by solving the α-shape of *in situ* immune substructures

Inflammatory dermatoses are characterized by the presence of large inflammatory infiltrates composed of specific immune cells and thought to be critical for the development of the pathology (e.g., type 2 immune cells and eosinophils in atopic dermatitis). To better understand the regional heterogeneity of pathological lesions from SLE, Kawasaki syndrome and COVID-toes, we took advantage of alpha (α)-shape algorithms that enable, by tuning the α parameter, to define a precise shape of sets of points by drawing bounding polygons based on the principle of Delaunay triangulation^43^. When combined with the digital immune landscapes generated with MANTIS, α-shape algorithms automatically generate polymorphic α-shapes around n-clusters composed of a minimum of 15 cells (**Fig. 4A**, i.e., 15 being the minimum number of cells often found in clusters of inflammatory but not in healthy-looking samples). This method enables to automatically detect and quantify the major inflammatory clusters (i.e., named thereafter “αROIs”) to provide a high-level view of the *in situ* immune architecture of the skin lesion for each patient and disease. We generated lymphoid (**Fig. 4B, C**) and myeloid (**Fig. 4D, E**) αROIs for all the samples. Healthy-looking controls displayed a few lymphoid αROIs, and 4/5 controls did not show myeloid αROIs. These data indicate that, in human skin at the steady-state, lymphoid cells have a tendency to form aggregates (i.e., composed of perivascular T lymphocytes^1^), while myeloid cells are more likely to be randomly distributed. In line with the data presented in **Fig. 3**, we found a higher proportion of both lymphoid and myeloid αROIs in pathological samples as compared to healthy-looking controls (**Fig. 4B-E**). Using global unsupervised hierarchical clustering of αROIs per disease, we can generate a high-level view of inflammatory clusters composition and observe trends in disease-specific immune responses (**Fig. 4F, G**). Notably, lymphoid αROIs of both Kawasaki syndrome and COVID-toes exhibited a significantly higher proportion of CD4^+^CD57^low^ T cells than that of SLE patients (**Fig. 4F, H**). Interestingly, both COVID-toes and SLE lesions displayed significant clusters of CD8^+^CD57^high^ cytotoxic T cells, highlighting the cytolytic aspect of pathological lesions microenvironment in these conditions (**Fig. 4F, I**). We next analyzed myeloid αROIs for all cases. We found that, COVID-toes had a particularly high density of clusters enriched in activated HLA-DR^high^ dDCs (**Fig. 4G,J**). Conversely, Kawasaki syndrome lesions showed an enrichment in both HLA-DR^high^ LCs (**Fig. 4G,K**) and mast cells (**Fig. 4G,L**), while SLE lesions displayed large aggregates of eosinophils (**Fig. 4G,M**). This finding is consistent with previous reports of strong eosinophilia in SLE^44–46^. No significant differences were observed regarding other immune cell subsets in αROIs (unpublished). While the precise role played by specific inflammatory clusters of immune cells in each disease remains elusive, these data strongly suggest that combining MANTIS digital maps with α-shape-based algorithms can reveal a significant non-random distribution of skin immune cells in skin lesions, with the presence of disease-specific immune structures. MANTIS analytic pipeline can thus enable to quickly solve the spatial organization of complex immune environments and open interesting perspectives for future investigations in the field of dermatopathology.

**Figure 4.**
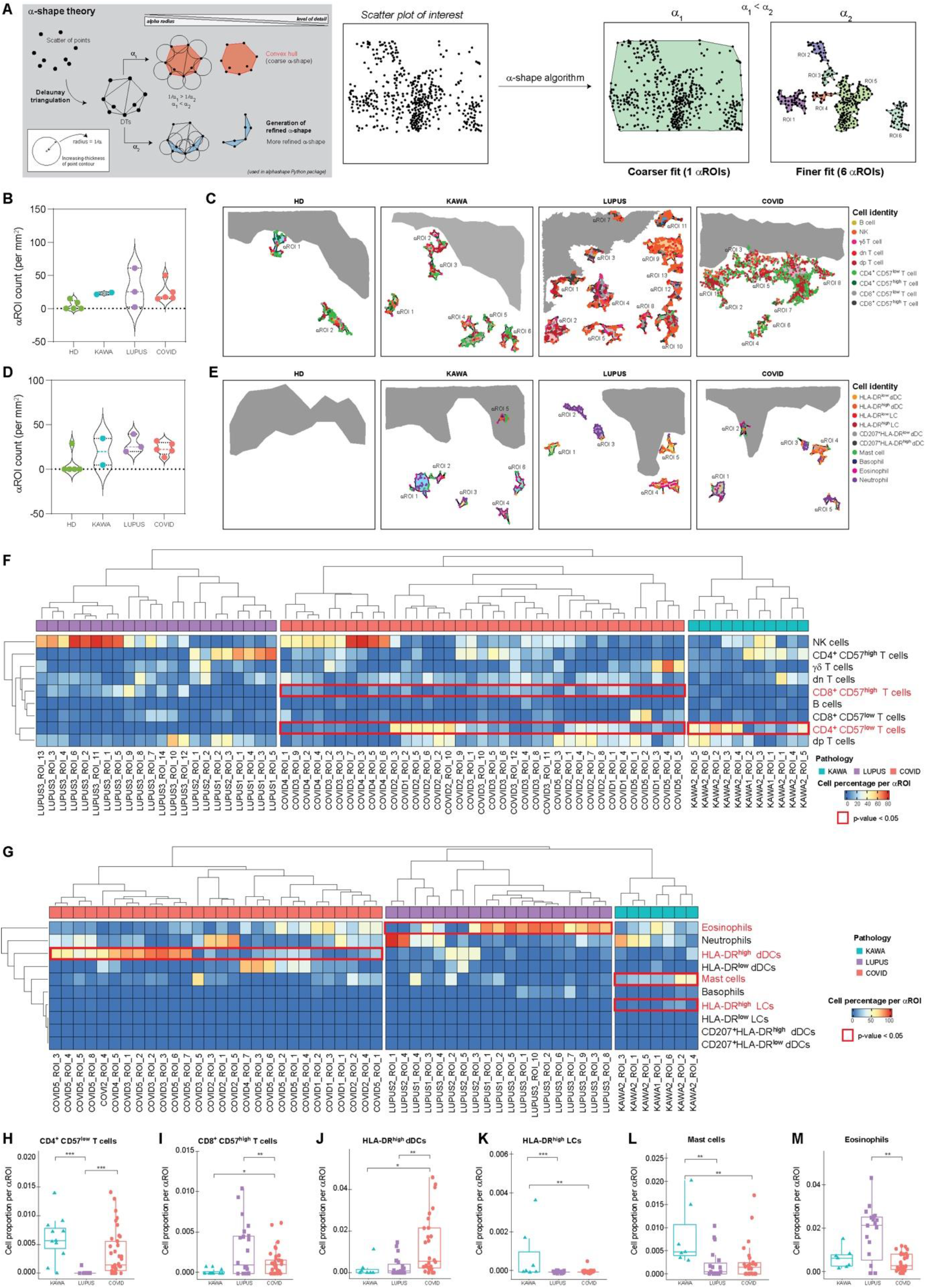
Automatic detection of α-regions of interest (αROI) enables exploration of inflammatory cluster topography in healthy and diseased skin. **A,** Alpha shape algorithm. Delauney triangulation of a given set of points formed a bounding polygon that contains all the points of the set. The alpha parameter was defined by the value α, and a circle with 1/α radius was drawn around each point of the dataset. The line between two circles meeting points formed a side of the bounding polygon, i.e., the alpha shape. α value defines the detail level of the alpha shape and allows modeling of voluminous structures (1/α_1_) or smaller structures (1/a_2_) having 1/α_1_ > 1/α_2_. **B, C,** Violin plot (B) and representative digital maps (C) of lymphoid α-ROI density in healthy and pathological skin. **D, E,** Violin plot (D) and representative digital maps (E) of myeloid α-ROI density in healthy and diseased skin. **F-K**, Mean proportion of CD4^+^ CD57^low^ T cells (F), CD8^+^ CD57^high^ T cells (G), HLA-DR^high^ dDCs (H), HLA-DR^high^ LCs (I), mast cells (J) and eosinophils (K) per αROI in diseased skin. Mean ± SEM; *P<0.05, **P<0.01, ***P<0.001 One-way ANOVA (F-K). **L, M,** Representative heatmaps of cell proportions in lymphoid (L) and myeloid (M) αROIs in pathological skin. A hierarchical clustering was applied on rows and on each pathology’s column.

## Discussion

Here we propose a general framework for 3-D quantitative and spatial analysis of skin immune cells at the cellular level. We first describe a simple method to perform a fast 3-D acquisition of up to 10 biomarkers simultaneously and extract a single-cell database containing the biological identity (including spatial coordinates) of skin lymphoid and myeloid cells. We then analyze the extracted databases using an automated and interactive analytic pipeline composed of phenotype attribution matrices coupled with cell-to-structure distance calculations and α-shape algorithm-based detection of major inflammatory clusters. Our analysis was focused on FFPE samples as it is still the most easily-available source of pathological tissues and can enable analysis of patients’ skin-sampled in routine clinical practice. However, the use of cryo-preserved samples is compatible with the approach we describe here, and enables the analysis of thicker tissue sections (unpublished data).

We identified that the first steps of the process, which consists in the generation of good quality 3-D multiplexed images with a significantly high ratio signal over autofluorescent background and no spectral spillover, were not necessarily obvious, while being critical for the rest of the study. This is why we emphasized the capability of a conventional 8-year-old (non-custom-built) confocal laser-scanning system to acquire 10 different fluorochromes simultaneously. This method of acquisition can be democratized to most academic/clinical facilities since they involve a conventional equipment coupled to basic spectral spillover compensation and single-cell data extraction strategies, via the use of commercially-available softwares (Huygens and Imaris; described in detail in the Methods section).

Single-cell segmentation is also very critical as it will constitute the very core of the future analysis of immune subpopulations and expression of biomarkers. Possible mistakes made at this step, e.g. the inability to separate immune cells in large infiltrates, would then result in misinterpretation of MANTIS-generated results. We tested different approaches to automatically segment healthy-looking and inflammatory skin samples, including random forest-based classifiers (e.g., Ilastik machine learning). While such a method was suitable to segment healthy-looking images with a low concentration of immune cells, it failed to segment complex inflammatory lesions, where large and packed immune clusters were present (unpublished). We thus opted for a semi-supervised segmentation of single immune cells using the software Imaris, in which the segmentation of each inflammatory cluster was quality-controlled manually and in 3-D (**Fig. 1F**). While this approach is probably more time-consuming, we could ensure a precise 3-D segmentation and further extraction of an accurate single-cell database to be processed with MANTIS. A recent study has reported the use of an analysis pipeline, including a new segmentation strategy, adapted from the field of astronomy named “AstroPath”^47^. Using this approach and only six biomarkers, they could identify important pathological features in biopsies from melanoma patients. These results, in line with our findings, highlight the importance of carefully selecting a list of biomarkers to be studied and of having the right analytic pipeline to draw reliable insights.

Because immunologists are more commonly used to identify immune cell populations with manual gating of populations based on flow cytometry, we validated the quantitative performance of MANTIS by analyzing the extracted single-cell databases for the 15 patients analyzed with the conventional flow cytometry software, FlowJo. We found that the MANTIS-based analysis on 3-D images generated with two different panels, and just a minimal number of 10 antibodies per panel, were sufficient to distinguish 19 immune subsets and identify disease-specific trends in skin lesions.

Because MANTIS attribution matrices can be quickly adjusted to any sets of markers, they could be compatible with single-cell databases generated using technology with high multiplexing capabilities such as CODEX^7,8^, MIBI-TOF^9^, IMC^10^, MuSIC^11^, CyCIF^12^, Cell Dive^13^ and others^14^. We included in MANTIS the α-shape algorithm that enables us to define the precise shape of inflammatory immune clusters based on the principle of Delaunay triangulation^43^. When applied to digital immune landscapes, the α-shape algorithm automatically identifies and quantifies dermal and epidermal inflammatory clusters (i.e., αROIs) composed of a minimum of 15 immune cells (**Fig. 4A**). This method enables one to directly analyze the major αROIs to provide a fast high-level view of the skin immune architecture in a given lesion. Using this method, we could quickly identify that immune subset (e.g., mast cells, HLA-DR^high^ dDCs, CD8^+^CD57^high^ cytotoxic T cells, eosinophils etc.) were specifically enriched in dermal areas only in some patient subgroups. These preliminary observations suggest that, depending on their etiology, pathological lesions could be due to distinct pathological mechanisms. This type of analysis opens interesting perspectives for the 3-D cartography of complex inflammatory skin lesions and should be pursued by additional studies on a larger number of patients. Importantly, while 2D immune landscapes are represented here to facilitate the visual assessment on Figures (3-D graphs are hardly perceivable on static pictures), the single-cell segmentation and extraction of cellular spatial coordinates were all performed in 3-D.

Based on CODEX high multiplexing capacity, previous studies^8,48^ have shown that it is possible to generate a high-level view of the cell-to-cell interaction landscape based on the principle of the Delaunay neighborhood graph^49^. The MANTIS α-shape algorithm is complementary, as it automatically identifies major immune structures while deciphering their cellular composition. Combining α-shape and neighborhood approaches could help to quickly solve the biology of major inflammatory clusters in the skin, by drawing the ligand-receptor interactome of immune and structural cells within the identified cluster. Such a high dimensional analysis of the skin immune architecture could provide a promising avenue for understanding the complexity of inflammatory skin manifestations with potential benefits for patient stratification and/or diagnosis.

There is a strong need to design new tools to assist clinical decision making and/or better apprehend the complexity of inflammatory dermatoses. While very promising processes have been made in the field of spatial biology^50–52^, there is an unmet need for a non-expensive and standardized multiplexed imaging analytical framework capable of automatically resolving the immune architecture of an inflamed skin. Here we show that the MANTIS analytical system is uniquely positioned to examine numerous questions in the fields of skin immuno-biology and should lay the foundation for a fast and automated analysis pipeline of relevant *in situ* inflammatory environments in both research and clinical facilities.

## Materials and Methods

### Human skin samples

Control normal skin biopsies were obtained from Genoskin SAS (https://www.genoskin.com/). Genoskin has obtained all legal authorizations necessary from the French Ministry of Higher Education, Research and Innovation (AC-2017-2897) and the Personal Protection Committee (2017-A01041-52). Skin biopsies from patients with lupus erythematosus were obtained from Toulouse University Hospital. Biopsies of COVID-toes were obtained from Toulouse, Reims and Lyon University Hospitals. Skin biopsies of multi-system inflammatory syndrome were obtained from Reims University Hospital.

### Skin section preparation, histology, and staining

Human skin samples were either frozen in optimal cutting temperature compounds (OCT, Tissue-Tek, unpublished) or formalin-fixed and paraffin embedded (FFPE). 10 μm FFPE-tissue sections were heated at 95°C for 20 minutes. Sections were subsequently immersed into Xylene for 30 minutes, washed in a graded series of ethanol (100%, 95%, 70%, 50% and 30%for 5 minutes each) and abundantly washed with distilled water. They were then treated using a heat-induced epitope retrieval method as previously described^53^.

FFPE-tissue sections were blocked and permeabilized with PBS 0.5 % (w/v)% BSA (Sigma-Aldrich), 0.3 % Triton X-100 (Merck) for 30 to 60 minutes at room temperature, then incubated with fluorophore-coupled antibodies or unconjugated antibodies overnight at 4°C in the dark. The sections were then washed three times in PBS 0.5 % (w/v)% BSA, 0.3 % Triton X-100 and incubated, if needed, with secondary antibodies in PBS 0.5 % (w/v)% BSA, 0.3 % Triton X-100 for 2 hours at room temperature in the dark. Finally, samples were treated with an autofluorescence quenching solution named TrueView (Vector Lab) for 5 minutes. The slides were mounted in Mowiol medium (Sigma-Aldrich) and sealed with a coverslip.

All conjugated and unconjugated antibodies used in this study were validated in single immunostainings of human skin and tonsils (unpublished), and are listed in **Supplementary Table 1**.

### Acquisition

512×512 pixel Z-Stack Images were acquired using an 8-year-old confocal microscope SP8 (Leica Microsystems) equipped with a HC PL APO CS2 with 40X NA 1.3 oil objective, a UV diode (405nm) and four lasers in visible range wavelengths (405, 488, 532, 552 and 635nm). The setup was made up of five detectors (three hybrid detectors with high quantum yield compared to classical photomultiplier (PMTs) detectors, and two PMTs). Mosaic sequential images were acquired using the between-stack configuration in order to simultaneously collect individual 7/8 channels and tiles before merging them to obtain one single image. Use of the between-stack configuration and the modulation of the detectors’ detection windows help to reduce the leaking of fluorophores. Finally, a digital zoom of 1.9 was applied during the acquisition and a mosaic multicolor image was obtained and exported into a .lif format. Detection windows and microscope configuration used in our study are listed in **Supplementary Table 2**.

### Image deconvolution and correction of spectral spillover

3-D mosaic images were then imported into Huygens SVI software, in order to correct the signal by applying deconvolution and crosstalk correction. Two deconvolution methods were used: the express deconvolution (theoretical and fast) or the deconvolution wizard (possibility to use experimental or theoretical parameters and to adjust the background value). Automatic crosstalk correction estimation was obtained and the coefficients were slightly adjusted manually, if needed, for optimal spillover correction.

### Segmentation

3-D mosaic images were imported into Imaris software to separate objects (cells) using a 3-D surface segmentation. Before creating the surface objects in Imaris, classical image processing was required. For instance, defining a threshold, adding a median filter, and/or normalizing the layers were sometimes applied in order to clean the background. Images were either cleaned using the CD45 surface objects or other channels by applying appropriate masks for each channel. Then, segmentation was applied on the CD45 channel surface. Statistics were exported into .csv format.

### Segmentation troubleshooting

In some cases, the surface creation parameters were not efficient to automatically obtain good object creation, or the module was not sensitive enough to detect low intensity objects. In this case, the creation of small objects was done manually and the threshold selection was also reduced. If the detected object was below 1 μm, a manual object unification with surrounding objects of same intensity was performed.

### Statistical data exportation

Statistical properties of each segmented object (cell) in the processed 3-D Imaris Multiplex image were automatically calculated. Object *Volume, Sphericity, Area, xyz Position* and *Mean Fluorescence Intensity* (*MFI*) in all channels were exported as a .csv table.

### FlowJo analysis and gating strategies

Identification and density assessment of immune cell subsets were analyzed using classical histo-cytometry^32^.

Immune cell populations were gated in FlowJo software as follows:

B cells: CD45^+^ CD20^+^
NK cells: CD45^+^ CD20^-^ CD3^-^ CD57^+^
CD4^+^ T cells: CD45^+^ CD20^-^ CD3^+^ TCRγδ^-^ CD4^+^ CD8^-^ CD57^low or high^
CD8^+^ T cells: CD45^+^ CD20^-^ CD3^+^ TCRγδ^-^ CD4^-^ CD8^+^ CD57^low or high^
γδ T cells: CD45^+^ CD20^-^ CD3^+^ TCRγδ^+^
dn T cells: CD45^+^ CD20^-^ CD3^+^ TCRγδ^-^ CD4^-^ CD8^-^
dp T cells: CD45^+^ CD20^-^ CD3^+^ TCRγδ^-^ CD4^+^ CD8^+^
Mast cells: CD45^+^ Tryptase^+^
DC: CD45^+^ CD1c^+^ CD207^-^ HLA-DR^low or high^
LC: CD45^+^ CD1c^-^ CD207^+^ HLA-DR^low or high^
DC CD207^+^: CD45^+^ CD1c^+^ CD207^+^ HLA-DR^low or high^
Neutrophils: CD45^+^ CD1c^-^ CD207^-^ Tryptase^-^ Siglec8^-^ MPO^+^
Eosinophils: CD45^+^ CD1c^-^ CD207^-^ MPO^-^ Tryptase^-^ Siglec8^+^ CD123^-^
Basophils: CD45^+^ CD1c^-^ CD207^-^ MPO^-^ Siglec8^+^ CD123^+^

## Tissue annotation

### Implementation of MANTIS reference panels

In order to enable cell identification, we built a binary table containing a literature-based theoretical signature of biomarkers expressed in each cell population identified by the used panel (naturally depending on the used set of antibodies), known as the reference attribution panel. If a cell population is positive for a marker, the value is set to 1, otherwise it is set to 0. If a cell population can be positive for a marker, there are two columns, one with the value set to 1, the other with the value set to 0 (i.e., γδ T cells can express CD4 or not). Two reference tables were implemented and designated by lymphoid and myeloid reference attribution matrices.

### Dynamic adaptation of reference matrices

Sample heterogeneity led to different acquisition parameters (laser power, gain, etc.). In order to standardize data processing, we scaled the reference tables and dynamically adapted, for each sample, the table values according to the MFI values. In practice, the value “1” in the binary table was replaced by the maximum MFI value acquired in the corresponding channel from the tested sample.

### Automatic cell type identification

To annotate the segmented objects, a correlation matrix between the MFI table and the adapted reference panel was generated by performing a pairwise Spearman’s Rank Correlation using the R software (2021). Each object was then phenotypically assigned to the cell type having the highest correlation coefficient. Objects with multiple highest correlation coefficients were assigned as “Other” cell types.

### Accuracy validation

The accuracy of MANTIS automatic cell identification was verified by comparing quantification results to classical histo-cytometry^32^. Briefly, linear regression of cell type density was computed between both attribution methods and regression coefficients were calculated. Regression coefficients ranging between 0.75 and 1 reflect MANTIS technique robustness.

### Activation status detection

MANTIS panels were designed to not only include discriminant markers for cell attribution but also non-discriminant and informative markers, for instance, activation markers. The cell populations of interest (CD4^+^ and CD8^+^ T cells in the lymphoid panel, DCs, LCs and CD207^+^ DCs in the myeloid panel) as well as the activation markers that reflect the activation status of these populations (CD57 in the lymphoid panel, and HLA-DR in the myeloid panel) were defined in the MANTIS algorithm. This latter automatically computes the MFI density curve associated with the activation markers within the selected populations. Subsequently, the MFI corresponding to the first peak of the density curve is defined as the MFI value above which the cell is considered positive for the activation marker.

### Alpha (α)-shape calculation

α-shape was calculated using the alphashape Python package. Briefly, Delauney triangulation of a given set of points formed a bounding polygon that contains all the points of the set. The α parameter was defined by the value α, and a circle with 1/α radius was drawn in such a way that two points of the dataset are located on the boundaries of the circle and the circle is empty. For each empty circle found, the line between the two points formed a side of the bounding polygon, i.e., the α shape. As α decreased, the alpha shape changed from a convex hull (e.g., epidermis α shape, α = 0.4) to a more tightly-fitting bounding box resulting in more refined alpha shapes (e.g., region of interest alpha shape [αROI], α = 0.1).

### Cell to structure distance calculation and nearest neighbor search

x-y coordinates of epidermis α-shape contours were stored using the k-dimensional tree method, which allows data ranking and structuration. Briefly, data points were classified based on nodes and branches space-partitioning, allowing a fast nearest neighbor calculation. For a given point (cell) of the dataset, the nearest neighbor in the epidermis alpha shape was found and the distance defined by r was calculated using the *scipy.spatial* Python package^41^. The distance of cells contained in the epidermis α shape was set to 0.

### Data clustering and αROI analysis

Regions of interest (αROI, i.e., inflammatory cell clusters) were identified using the α shape algorithm with a tuned α parameter (α = 0.1), allowing correct detection of high cell density areas. αROI with less than 15 cells were removed from the analysis. For each selected αROI, specific characteristics were calculated and extracted, such as area, total number of cells, cell number, proportion by cell type and αROI center coordinates.

### Data visualization

Visualization charts were obtained using the *ggplot2, Pigengene* & *ComplexHeatmap* R packages, and *matplotlib & seaborn* Python packages. t-Distributed Stochastic Neighbor Embedding (tSNE) was computed with Rtsne.

### Statistics

Statistical tests were performed using Prism 8 (GraphPad Software), the *Rstats* and *rstatix* R packages. One-way ANOVA with Tukey’s test for multiple comparisons or Mann-Whitney test were performed on samples as noted in the respective figure legends. A p-value of less than 0.05 was considered statistically significant.

## Supporting information

Supplementary manuscript

## Authors’ contribution

N.G. conceived the project. M.S., R.H., J.M., N.S., M.T. and N.G. were involved in experimental design. M.S., R.H., J.M., N.S. and N.G. performed most experiments and compiled the data. M.T., E.P. and E.B. provided important help with experiments. M.T., L.L., D.J., B.C., C.L., C.B.L. and J.K. provided clinical samples and expertise. All authors participated in analyzing the data and writing or editing the paper. We thank all members of the Gaudenzio laboratory at Infinity and of Genoskin for discussions and technical assistance. We thank Sophie Allart and Simon Lachambre for technical assistance at the cellular imaging facility of Inserm UMR 1291, Toulouse. We thank Samantha Milia and Timothé Durand-Plavis for technical assistance at the experimental histopathology platform US06/CREFRE.

