## Supplementary manuscript for "3-D deconvolution of human skin immune architecture with Multiplex Annotated Tissue Imaging System (MANTIS)"

**Title**

### Supplementary Figures and legends

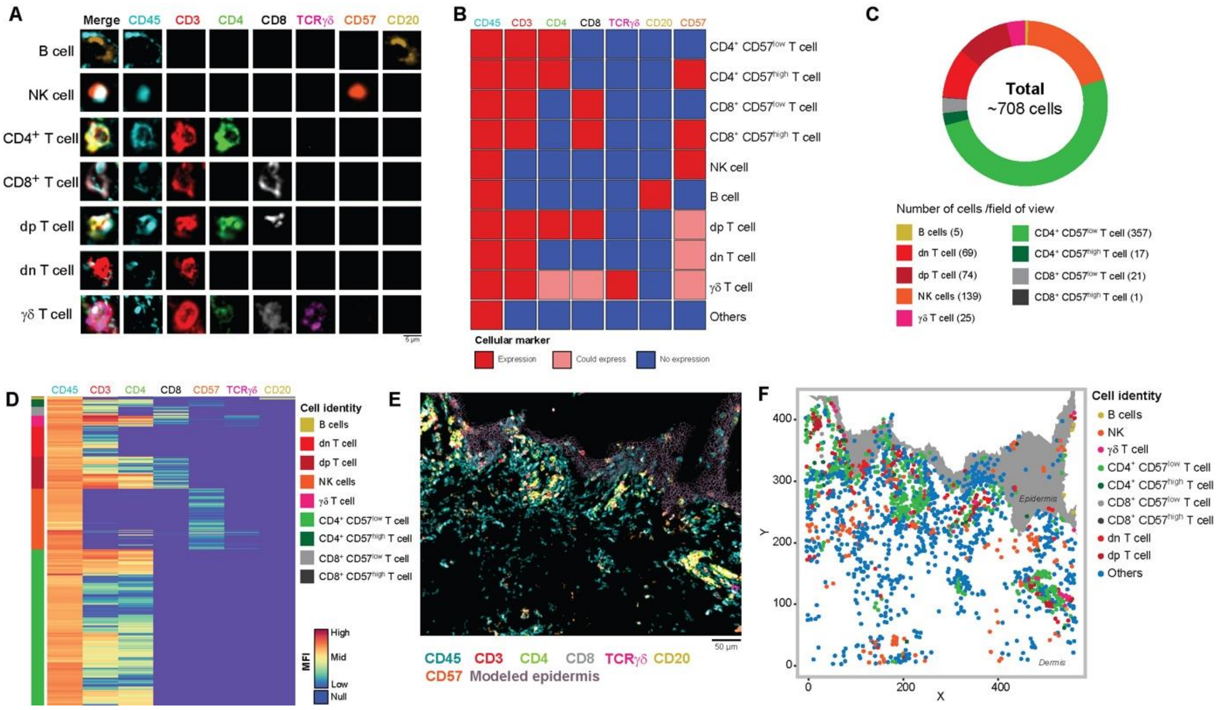


**Supplementary Figure 1. Main lymphoid cell populations annotated using MANTIS algorithm. A**, Examples of single-cell staining of all used biomarkers in identified lymphoid cells. **B**, MANTIS attribution matrix for the lymphoid panel. **C**, Tissue annotation and cell proportion of diseased skin (SLE patient). **D**, Heatmap of mean fluorescence intensity levels of used markers in identified lymphoid cells (colored scale). **E, F,** Representative 3-D confocal multiplex image (E) and associated digital map generated with MANTIS (F) of the lymphoid panel in diseased (SLE) skin. Scale bar = 50 µm.

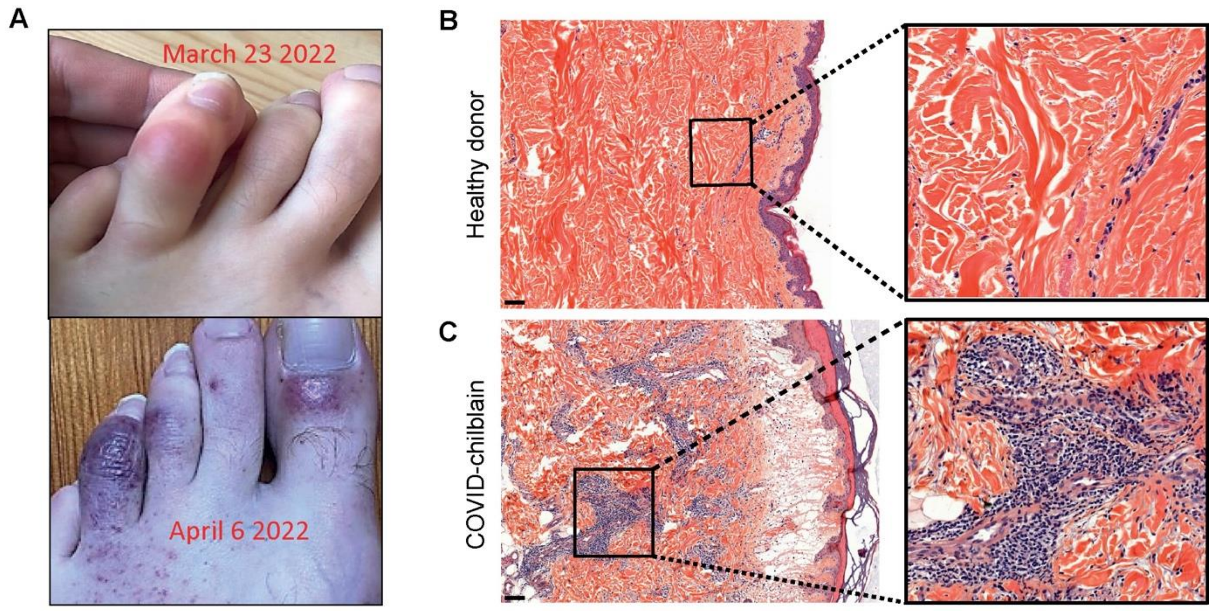


**Supplementary Figure 2. COVID-toes and associated histology. A,** Evolution of a chilblain-like lesion in a COVID patient between March 2022 (upper panel) and April 2022 (lower panel). **B, C**, Comparison of H&E staining of a healthy-looking skin sample (B), and a chilblain-like lesion associated with COVID (COVID-T=toes), showing immune infiltrates (C). Scale bar: 100µm.

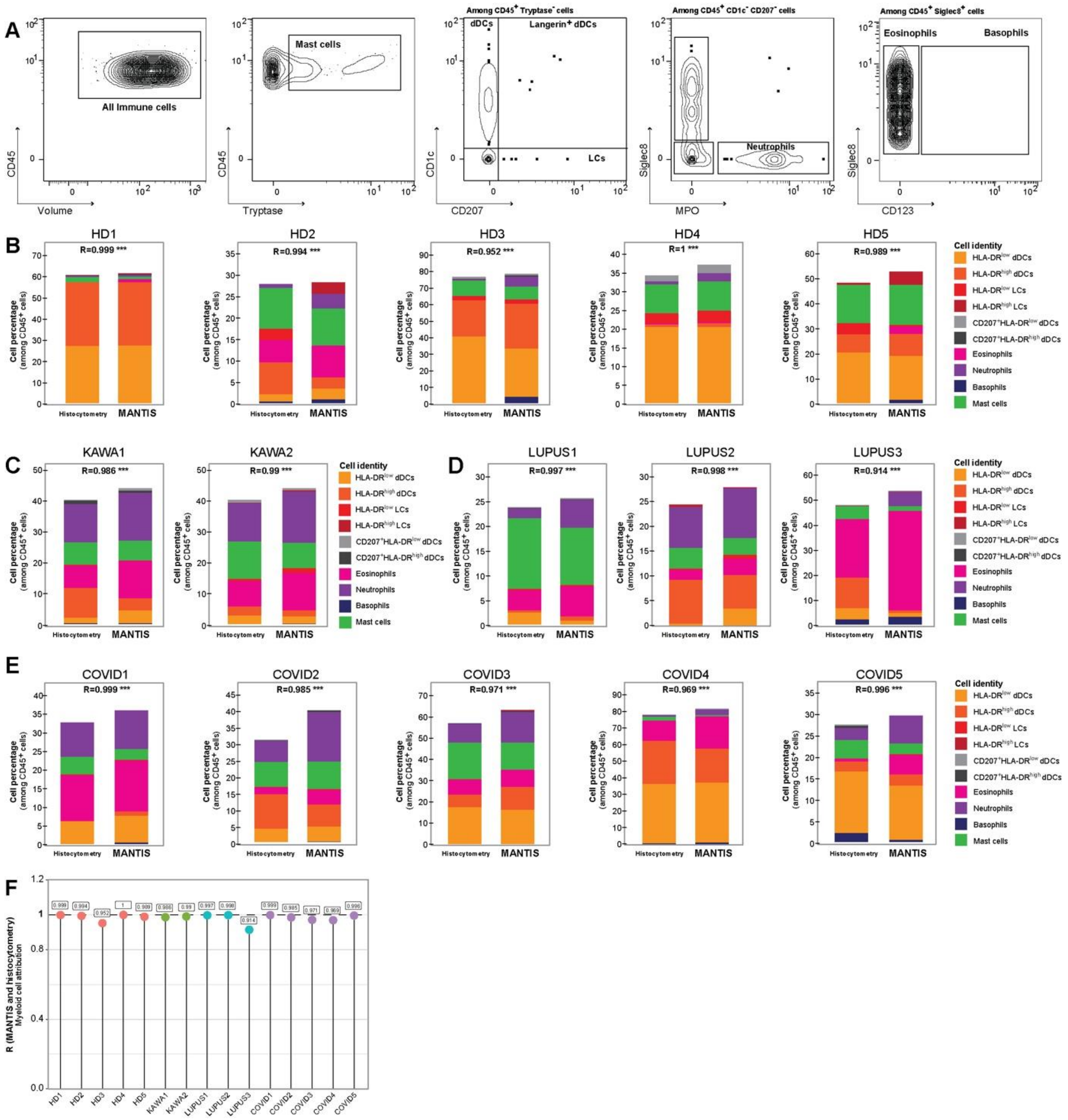


**Supplementary Figure 3. Quantitative validation of MANTIS myeloid tissue annotation with classical histo-cytometry. A,** Gating strategies of identified myeloid cell populations using FlowJo. **B-E**, Comparison of cell type percentages using classical histo-cytometry or MANTIS and associated Pearson correlation coefficient in healthy skin (**B**) and Kawasaki (**C**), SLE (**D**) and COVID-toes (**E**) lesions. ***P<0.001, Pearson correlation test. **F**, Lollipop chart of Pearson correlation coefficients of all samples comparing histo-cytometry and MANTIS cell attribution.

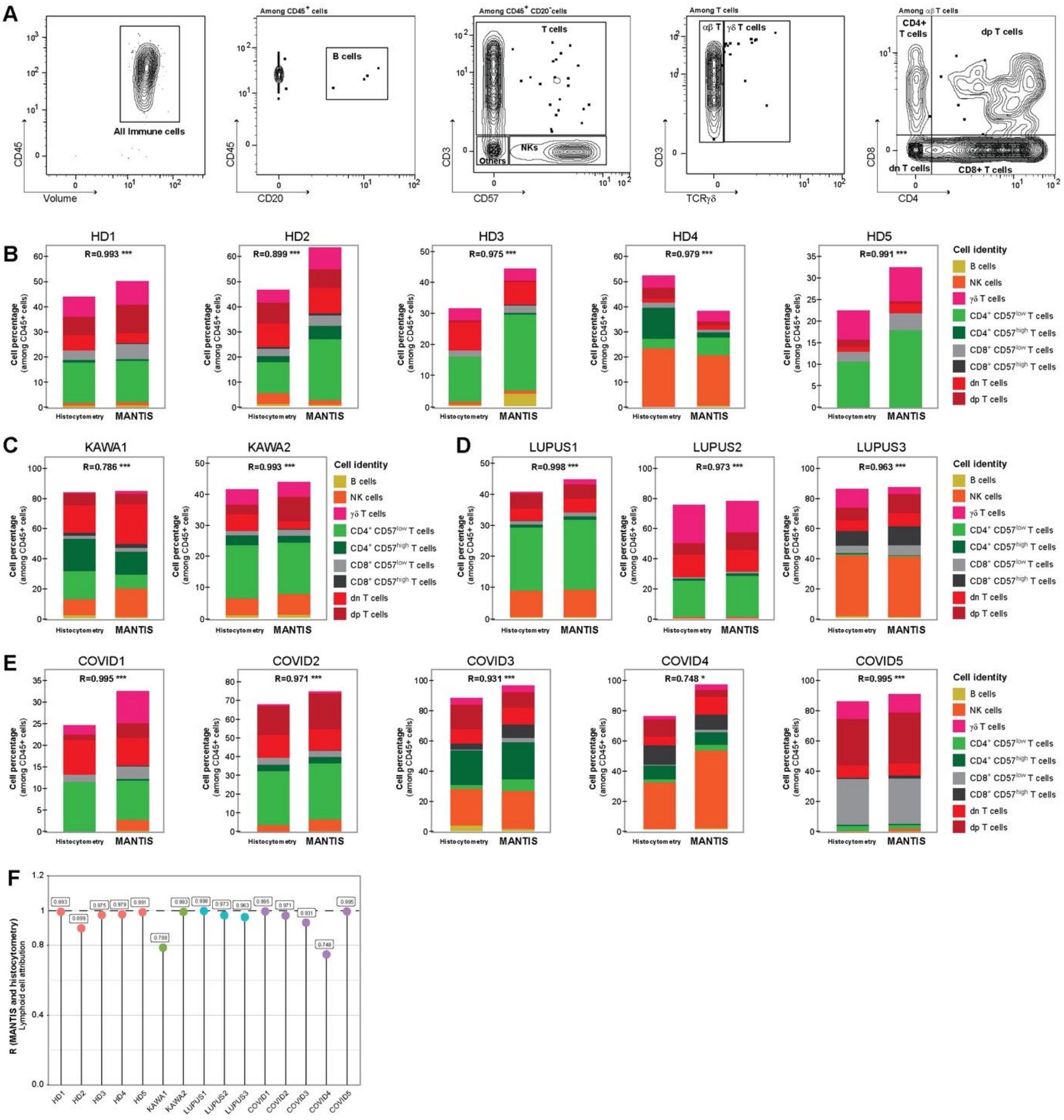


**Supplementary Figure 4. Quantitative validation of MANTIS lymphoid tissue annotation with classical histo-cytometry. A,** Gating strategies of identified lymphoid cell populations using FlowJo. **B-E**, Comparison of cell type percentages using classical histo-cytometry or MANTIS and associated Pearson correlation coefficient in healthy skin (B) and Kawasaki (C), SLE (D) and COVID-toes (E) lesions. *P<0.05 ***P<0.001, Pearson correlation test. **F**, Lollipop chart of Pearson correlation coefficients of all samples comparing histo-cytometry and MANTIS cell attribution.

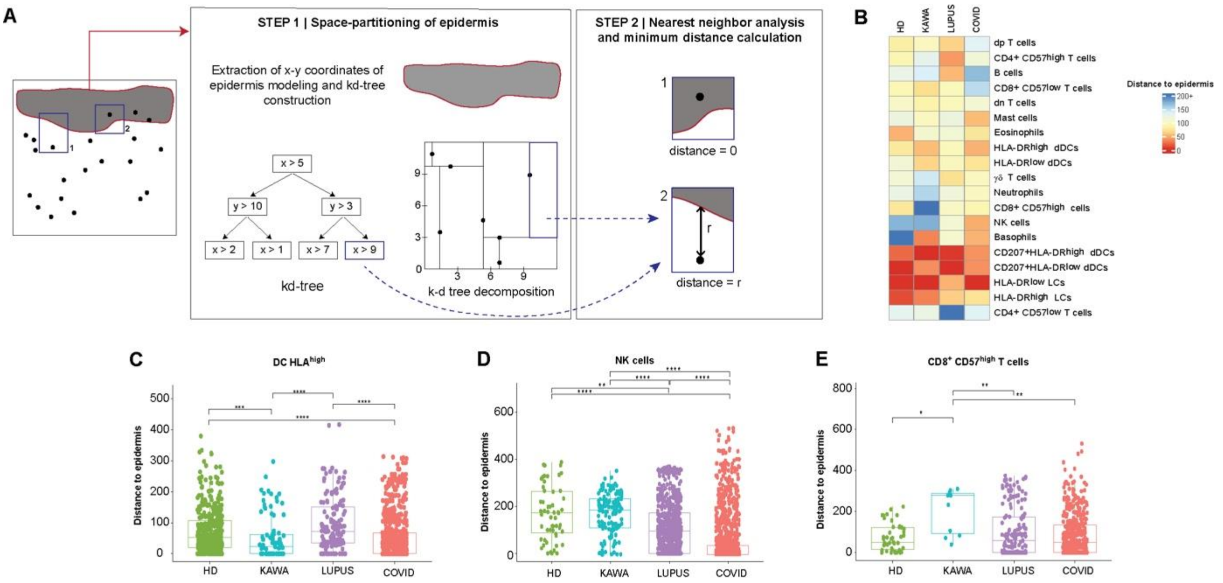


**Supplementary Figure 5. 3-D spatial distribution of immune cells and structural elements is computed using MANTIS. A**, Quick nearest neighbor computation using KD-tree space-partitioning. *xy* coordinates of the epidermis were stored based on tree decision, using KD-Tree. Nearest neighbor search using KD-Tree point storage was then computed for each detected cell, and the minimal distance was calculated. **B**, Heatmap of mean distance to epidermis (colored scale) per cell type in healthy and pathological skin. **C-E**, Mean distance to epidermis (in µm) of dDC HLA$^{high}$ (C), NK cells (D) and CD8$^{+}$ CD57$^{high}$ T cells (E) in healthy and pathological skin. Mean ± SEM; *P<0.05, **P<0.01, ***P<0.001 One-way ANOVA.

### Supplementary Tables and Legends

| Antibody | Clone | Fluorochrome | Concent. | Reference |
|---|---|---|---|---|
| CD45 | HI30 | AF-532 | 0.12 µg | Thermofisher, #58-0459-42 |
| **Lymphoid panel** | | | | |
| CD4 | N1UG0 | Unconjugated<br>*Revealed with anti-mouse AF488* | 10 µg/ml | eBioscience™, #14-2444-82 |
| CD3 | Polyclonal | Unconjugated<br>*Revealed with anti-rabbit AF594* | 10 µg/ml | Agilent, #A045229-2 |

| | | | | |
|---|---|---|---|---|
| CD20 | 2H7 | APC-Cy7 | 20 µg/ml | Biolegend, #302313 |
| TCR-γδ | B1 | BV650 | 15 µg/ml | BD, #564156 |
| CD8a | AMC908 | eFluor 660 | 10 µg/ml | Thermofisher, #50-0008-80 |
| CD57 | TB01 | eFluor 450 | 10 µg/ml | Thermofisher, #48-0577-41 |
| **Myeloid panel** | | | | |
| Siglec8 | Polyclonal | eFluor 450<br>*Revealed with anti-rabbit AF405* | 1:50 | Thermofisher, #PA5-110774 |
| CD1c | L161 | Zenon AF488 Using Zenon Mouse IgG1 labeling kit (Thermofisher, #Z25000) | 10 µg/ml | Biolegend, #331502 |
| Tryptase | AA1 | Zenon AF647<br>Using Zenon Mouse IgG1 labeling kit (Thermofisher, #Z25000) | 1:50000 | Abcam, #ab2378 |
| CD207 | 923B7 | AF546 | 5µg/ml | Biotechne, #DDX0373A546 |
| MPO | Polyclonal | Unconjugated<br>*Revealed with anti-goat-AF594* | 5 µg/ml | Biotechne, #AF3667 |
| HLA | L243 | AF700 | 10 µg/ml | Biotechne, #NB100-77855AF700 |
| CD123 | 6H6 | SB645 | 1 µg | eBioscience™, #64-1239-42 |

**Supplementary Table 1. Key resources.** Classical and available unconjugated and conjugated antibodies were used in this study at indicated concentrations.

| Antibody | Fluorochrome | Excitation laser (nm) | Laser power (%) | Detector type | Detection window |
|---|---|---|---|---|---|
| CD45 | AF-532 | 532 | 6 | HyD | 535-578 nm |
| | **Lymphoid panel** | | | | |

| | | | | | |
|---|---|---|---|---|---|
| CD4 | AF488 | 488 | 2.7 | HyD | 503-539 nm |
| CD3 | AF594 | 552 | 6 | HyD | 603-633 nm |
| CD20 | APC-Cy7 | 635 | 50 | PMT | 740-790 nm |
| TCR-gd | BV650 | 405 | 10 | HyD | 624-682 nm |
| CD8a | eFluor 660 | 635 | 67.2 | HyD | 651-693 nm |
| CD57 | eFluor 450 | 405 | 10 | HyD | 415-479 nm |
| | **Myeloid panel** | | | | |
| Siglec8 | eFluor 450 | 405 | 3.67 | HyD | 409-470 nm |
| CD1c | AF488 | 488 | 1 | HyD | 505-536 nm |
| Tryptase | AF647 | 635 | 89.5 | HyD | 648-689 nm |
| CD207 | AF546 | 552 | 4 | HyD | 558-588 nm |
| MPO | AF594 | 552 | 2.04 | HyD | 608-633 nm |
| HLA | AF700 | 635 | 50 | PMT | 699-782 nm |
| CD123 | SB645 | 405 | 3.97 | HyD | 625-681 nm |

**Supplementary Table 2. Microscope configuration of MANTIS acquisitions.** Between-stack acquisition parameters were configurated using all available four lasers in visible range wavelengths and detectors (Hybrid [HyD] or photomultiplier [PMT]).
